# In silico evidence of superantigenic features in ORF8 protein from COVID-19

**DOI:** 10.1101/2021.12.14.472240

**Authors:** Guillermo Gómez-Icazbalceta, Zubair Hussain, Marcela Vélez-Alavez

**Author notes:** Correspondence: Dr. Guillermo Gómez-Icazbalceta.

## Abstract

Very early on COVID-19 pandemic outbreak, it was noted that the some of the virus-induced clinical conditions resembled features of toxaemia caused by the toxic shock syndrome toxin type 1, which is a soluble superantigen produced by *Staphylococcus aureus*. Among all SARS proteins, the ORF8 protein from SARS-2 virus is significantly different from other known SARS-like coronaviruses, and therefore could exhibit unique pathogenic properties. We assess if ORF8 protein bears super antigenic features using *in silico* tools. We show that ORF8 has properties of an extracellular soluble protein and shares a significant degree of amino acid sequence identity with toxic shock syndrome toxin. Besides, docking and binding affinity analyses between monomeric and homodimeric ORF-8 with Vβ 2.1 and TRBV11-2 reveal strong interaction and high binding affinity. ORF8-TRBV11-2 strong interaction can contribute to the observed clonal expansion of that chain during COVID-19-associated multisystem inflammatory syndrome. Taken together, the evidence presented here supports the hypothesis that ORF8 protein from SARS-2 bears super antigenic properties.

## INTRODUCTION

It is well established that SARS-2 virus actively replicates in the lungs causing severe pneumonia [1], along with other related pathologies, whose manifestations differ among age groups and comorbidities presented during infection [2]. However, extrapulmonary conditions overlapping with some features of superantigen-induced toxic shock syndrome (TSS) are also frequently expressed, such as systemic hyper inflammation and coagulation disorders [3-5]. TSS is caused by the toxic shock syndrome toxin type 1 (TSST), a soluble superantigen (Sag) produced by *Staphylococcus aureus* [6]. Systemic inflammatory condition during COVID-19 infection, properly referred to as cytokine release syndrome (CRS) [7], is characterized by the release of elevated levels of inflammatory cytokines, such as IFN-γ, IL-1β, IL-10, IL-6, IL-8, TNFα, MCP-1, and GM-CSF, that induce excessive systemic thromboinflammation, and eventually may lead to disseminated intravascular hypercoagulation, multi-organic failure and a high risk of death. Besides, CRS induced by COVID-19 infection triggers a plethora of systemic immune disorders. More than 50 immune disorders have been described so far [8]. In this regard, a condition referred to as multisystem inflammatory syndrome (MIS), characterized by an immunopathophysiology resembling bacterial toxic shock syndrome has been observed during advanced COVID-19 infection, mainly in pediatric population [9, 10].

During MIS in children (MIS-C), a clonal expansion of the T cell receptor (TCR) β variable gene 11-2 (TRBV11-2) also known as Vβ21.3, is observed, representing up to 24% of the total T cell clonal space [11, 12]. The massive expansion of a single T cell clone is a key feature of bacterial Sag TSST [13]. Also, a cascade of inflammatory cytokines and thrombotic mediators are released, leading to systemic thrombotic inflammation [12]. Taken together, these lines of research indicate that MIS-C has many characteristics of TSS. An *in silico* research has elucidated a potential superantigen located on the spike protein of COVID-19 virus, that could be able to interact directly with human TCR [14]. Such interaction was discovered to be driven by a peptide high related to enterotoxin B Sag.

However, cumulative evidence suggests that thromboimmune exacerbation can be also driven by soluble viral factors acting independently of viral core [15-17]. We aimed then to discover nonstructural viral proteins that could bear Sag features.

Among all SARS proteins, the open reading frame 8 protein (ORF8) is significantly different from other known SARS-like coronaviruses, therefore could provide unique pathogenic properties [18]. ORF8 is a 121 amino acids nonstructural protein [19] and it is one of the most rapidly evolving βcoronavirus proteins [20]. Its only available crystal structure showed a homodimeric protein [20], and monomeric protein was also eluted from the cell extract. In contrast, its function has been barely characterized. ORF8 can directly interact with major histocompatibility complex I (MHC-I) and down regulate their expression on different cell types, probably resulting on disruptions of the adaptive immune system [21]. ORF8 could also participate as a molecular host factor of mimicry in immune evasion strategy [22]. Regarding immune activation, ORF8 can trigger a cytokine storm in a mice model, indicating it has an important role in the pathogenesis of CRS [23].

Therefore, we undertook an in silico approach to estimate the Sag properties of this protein. Our research shows that ORF8 from SARS-2 virus bears properties of soluble and extracellular protein, shares a high degree of similitude with bacterial TSST, and forms a strong docking with β chains 2.1 and 21 of human TCR. where the latter is highly expanded during COVID-19-associated multisystem inflammatory syndrome.

## METHODOLGY

### Sequences

All the protein FASTA sequences were retrieve from NCBI protein databank. ORF8 with accession number YP_009724396.1; TSST with accession number WP_061822158.1.

### *In silico* analyses of ORF8 basic properties

Basic properties were predicted using protein FASTA sequence as input. Subcellular localization of ORF8 was predicted using Deeploc 1.0, a deep learning algorithm software that predict protein localization among 10 different cell compartments [24]. The signal peptide in ORF8 was predicted using SignalP 5, an algorithm software that predicts signal peptides and cleavage site [25]. Basic secondary structure of ORF8 was predicted using PredictProtein, an algorithm software that predict secondary structure of proteins based on a wide array of protein features [26].

### Amino acid similarity analysis

The amino acid (aa) similarity analyses between ORF8 and TSST were performed using protein BLAST, [27] and Clustal Omega [28], with default settings.

### Models retrieval

Crystal structures were retrieved from Protein data bank [29]. ORF8 homodimer was retrieved from PDB with ID 7JTL. Vβ 2.1 was retrieved from PDB with ID 3MFG and TRBV11-2 was retrieved from PDB with ID 6R2L. All models were purified by removing water molecules and ligands using PyMol version 2.4.1 [30]. Chain B and Chain E in 3MFG and 6R2L were Vβ 2.1 and TRBV11-2, respectively, so the rest of chains were deleted using UCSF Chimera version 1.14 [31] Missing residues in Vβ 2.1 were added by using user template of Swiss model, followed by energy minimization using Swiss PDB Viewer version 4.1.0 [32].

### Modelling of monomeric ORF8

Model of monomeric ORF8 was developed using Galaxy TBM (33) [33], followed by energy minimization using Swiss PDB Viewer version 4.1.0 and refinement using Galaxy refine tool of Galaxy web [34]. Verification of model was carried out using VERIFY3D and ERRAT [36]. Ramachandran analyses were carried out using ProCheck [37].

### Binding sites determination

Binding sites determination for Homodimer ORF8, Vβ 2.1, TRBV11-2 and monomeric ORF8 was carried using SPPIDER [38] and Meta-PPISP [39] and common binding site residues from both servers were used as binding sites residues for docking.

### Protein-protein docking

Docking of homodimer ORF8, and monomeric ORF8 with Vβ 2.1 and TRBV11-2 was carried out using Cluspro [40]. Best docked complexes were selected based on maximum member size and lowest energy from ClusPro. The interacting amino acids residues analysis were done using PyMol version 2.4.1 [30].

### Binding affinity

Binding affinity for the docked protein-protein complexes obtained from ClusPro were carried out at 25°c by Prodigy [41]. 3MFG was included as a reference. The 3MFG were given to server after removing water molecules and ligands and co-factors by using PyMol version 2.4.1 [30].

## RESULTS

### 1. Signal peptide

An N-terminal signal peptide for COVID-19 ORF8 accessory protein was predicted with high probability by SignalP5 software. The cleavage site was estimated at position 15 (Table 1 and Figure 1).

**Table 1.** Predicted signal peptide values for ORF8 by SignalP5.

| Protein type | Likelihood | Other |
| --- | --- | --- |
| Signal peptide | 0.9973 | 0.0027 |

**Figure 1.**
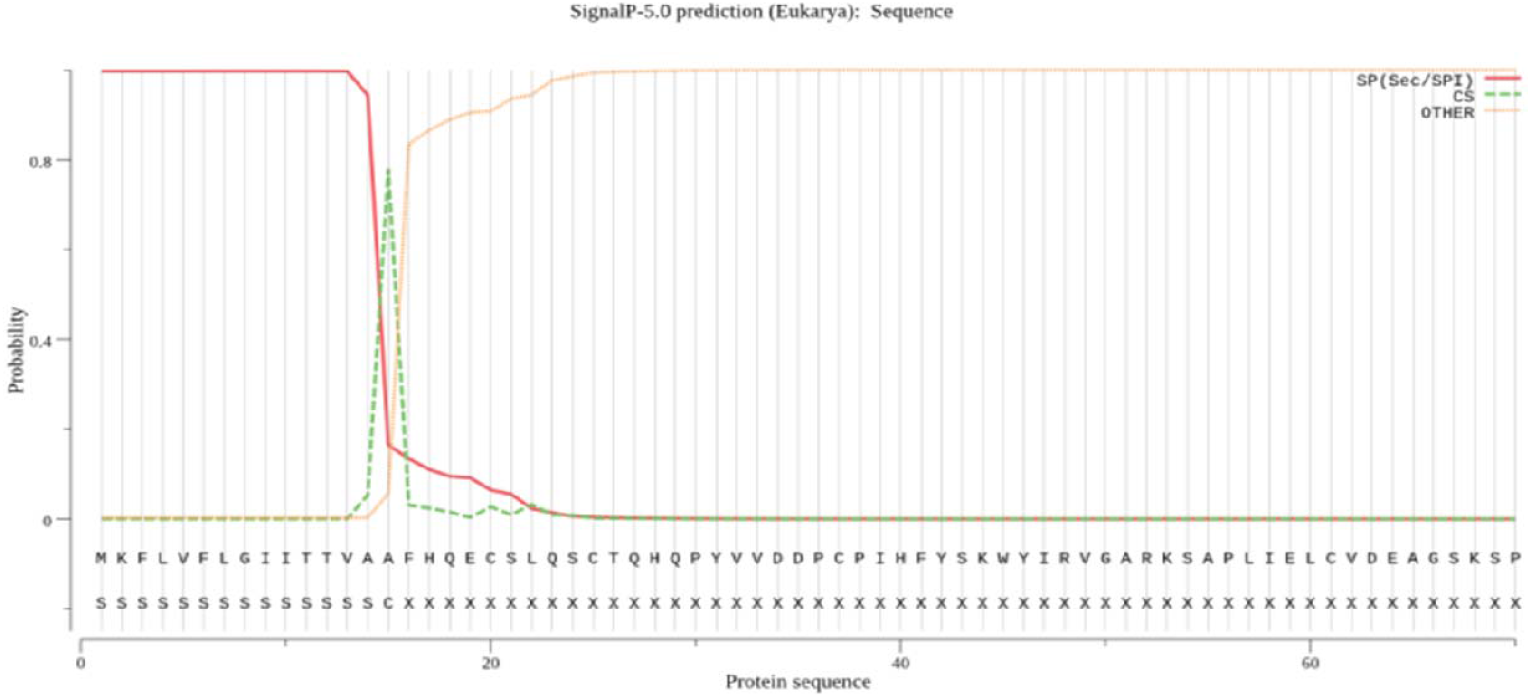
Probable location of predicted signal peptide in ORF8. Red line: predicted signal peptide; dashed green line: location of cleavage site; orange line: predicted for other than signal peptide.

### 2. ORF8 Protein destiny

The subcellular location of ORF8 was predicted using DeepLoc software. The software predicted ORF8 as an entirely soluble and extracellular protein and bearing a N-terminal signal peptide (Figure 2).

**Figure 2.**
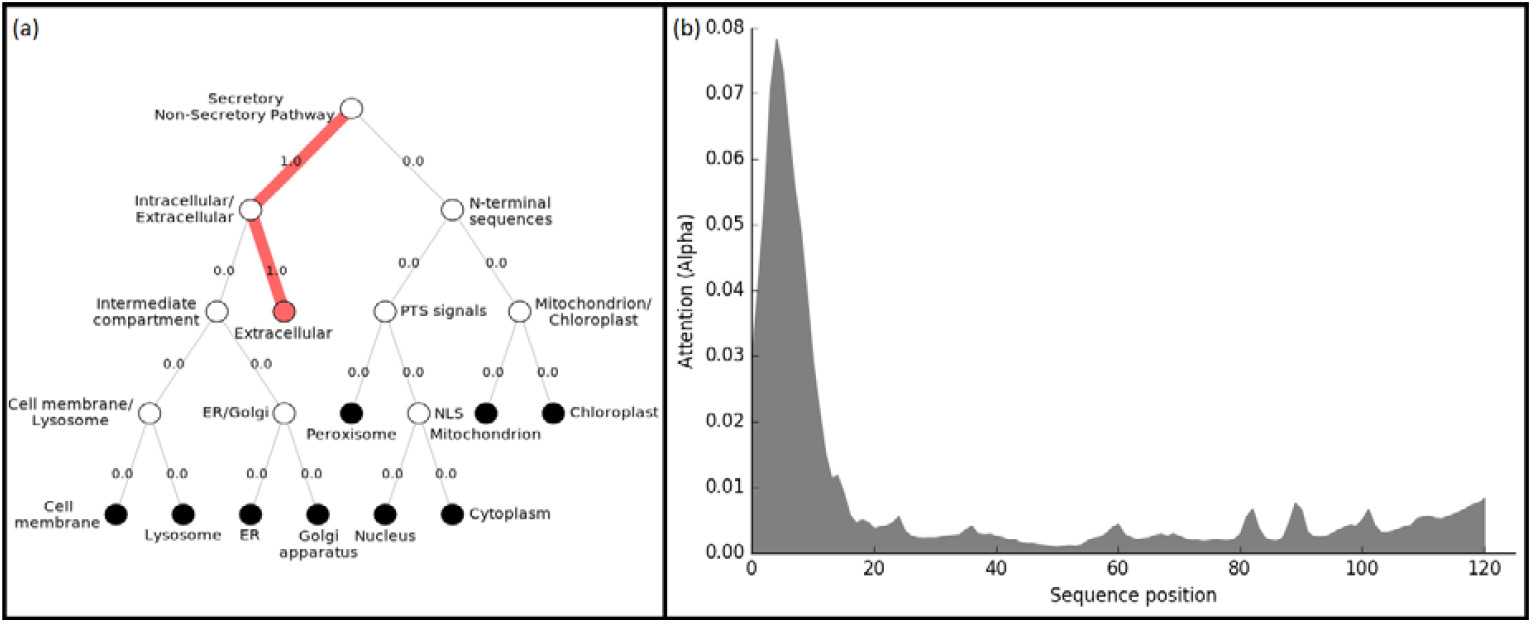
ORF8 was predicted to be entirely a soluble and extracellular protein, with the N-terminal region as relevant for these features, according to analysis with the DeepLoc-1. **a)** Hierarchical tree shows in red lines the predicted subcellular localization of ORF8. **b)** Histogram shows N-terminal region predicted as the most relevant zone related to those features, according to Attention score value

The predicted probability of ORF8 for bearing a signal peptide at N-terminal region was nearly 1, while for the rest of the peptide it was zero. The cleavage site was predicted at position 15. Therefore, the N-terminal region is the only region of ORF8 with a putative signal peptide, which is a feature of extracellular proteins. The software also predicted ORF8 as soluble and extracellular protein and gave to each of these properties a score of 1, which is the maximum score. Therefore, ORF8 is predicted to bear properties of a soluble circulating protein.

### 3. Predicted secondary structure

To further determine the properties of ORF8 protein, the secondary structure of ORF8 was predicted using PredictProtein platform (Table 2).

**Table 2.** Description of predicted secondary structure of ORF8 by PredictProtein

| Helix | Strand | Loop | Disulfide bridges |
| --- | --- | --- | --- |
| 1.89% | 40.57% | 57.55% | aa 68 to 87; length 20 aa |

The platform predicted ORF8 is constituted by a loop in near 60% of its length. Loops are associated to ligand-receptor interactions and therefore the predicted loop could serve ORF8 to interact with other proteins. Notably, a disulfide bond of 20 aa length was also predicted, spanning from amino acids 68 to 87. The presence of a disulfide bond is a feature of soluble proteins, as it provides resistance to degradation and hence improves their half-lifes during exposure to extracellular milieu [42].

### 4. ORF8 and bacterial toxins similitude analysis

To determine if ORF8 shares a degree of sequence similitude with TSST from *Staphylococcus aureus*, we ran a BLAST analysis using ORF8 as a query. We found that TSST shared 34.29% of amino acid identity with ORF8 (E = 0.049), with a query coverage of 60%, in 3 alignments (Figures 3 and 4). The shortest alignment, comprising N-terminal region, may belong to signal peptide. The other two alignments spanned aa 32 to 91 of ORF8 and thus may be more related to a superantigen-like function.

**Figure 3.**
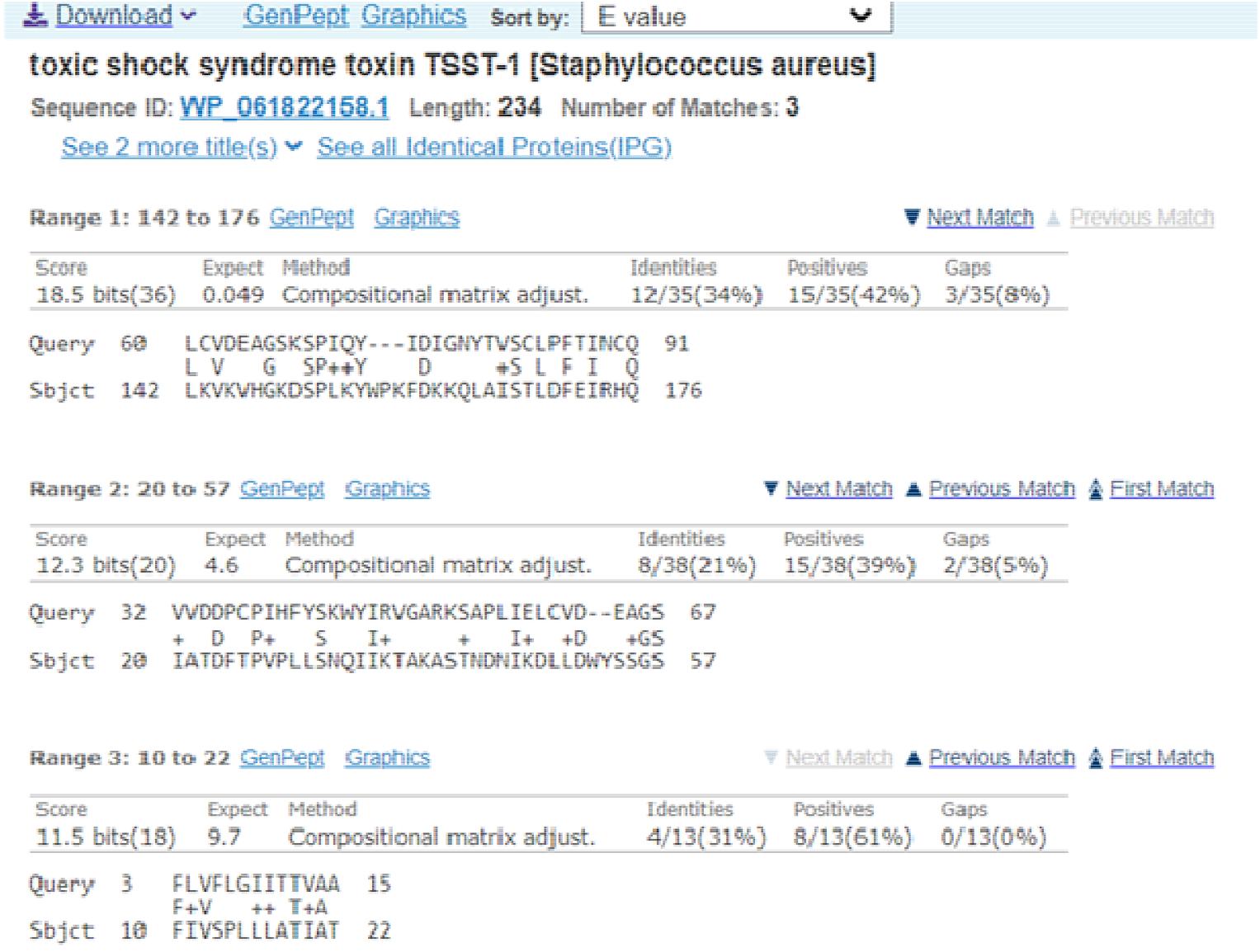
Screenshot of alignment between ORF8 from COVID-19 (Query) and TSST from SA bacteria (Subject), according to NCBI Blast analysis.

**Figure 4.**
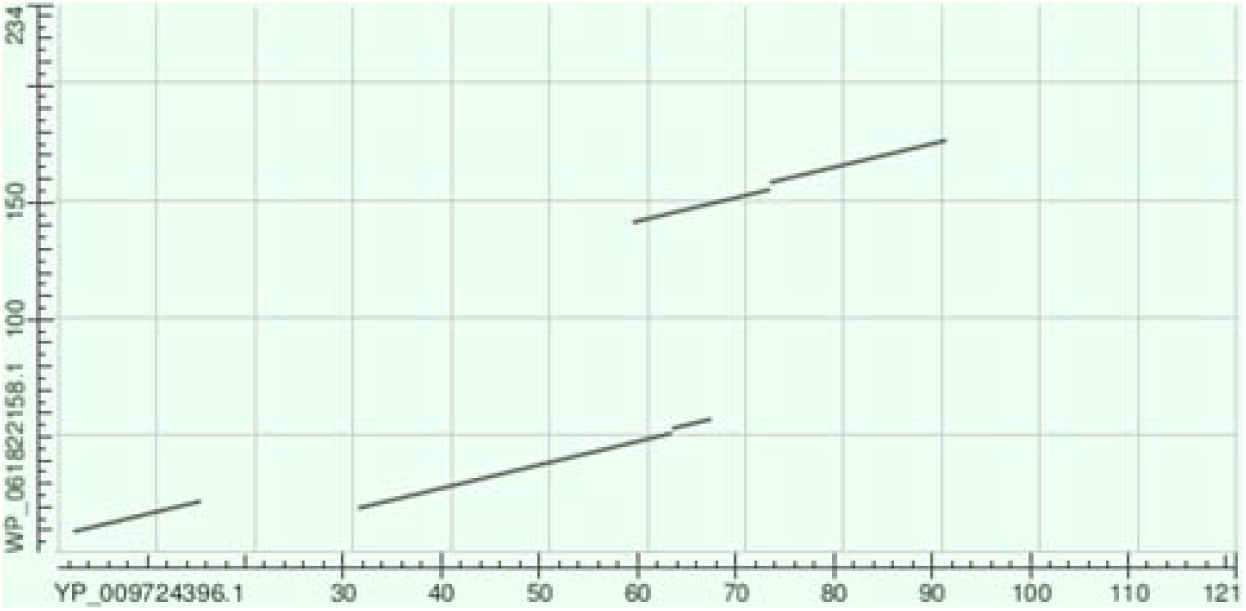
Screenshot of dot matrix plot of amino acid sequences similarity between ORF8 (X axis) and TSST from SA (Y axis), according to NCBI Blast analysis.

### 5. Clustal similitude analysis

To obtain additional support to the BLAST findings, we ran a similitude analysis in Clustal Omega. The analysis showed that ORF8 had significant degree of similarity with TSST in a region spanning aa 60 to 95 of ORF8, where 24 positive matches were identified. Notably, in the same region, aa 91 and 94 of ORF8 matched aa Q136 and Q139 from TSST, which have been defined as essentials for this bacterial superantigen to TRBV11-2 chain docking [43, 44]. Thus, ORF8 had a significant region of similarity to TSST in a region essential for docking to Vβ 2.1

### 6. Modelling, refinement, and validation of monomeric ORF8

Predicted model of monomeric ORF8 by Galaxy TBM were subjected to refinement by Galaxy refine tool after minimization by SWISS PDB viewer. Refinement of monomeric ORF8 by Galaxy refinement showed 5 models, out of which the model number 1 was selected as the best model. Comparison of model generated for monomeric ORF8 by Galaxy TBM and Galaxy refine tool is shown in Table 3. Then, the model quality was evaluated by Verify 3D, ERRAT, and Pro Check. Verify 3D program was used for verification of the correctness of the model (Figure 6). The verified 3D score for ORF8 was 82.64%. Reliability of monomeric ORF8 model were evaluated by ERRAT (Figure 7). ERRAT score for the monomeric ORF8 was 83.3%.

**Table 3.** Galaxy TBM and Galaxy refine scores for modelled monomeric ORF8.

| Model | GDT-HA | RMSD | Mol Probity | Clash score | Poor rotamers | Rama favored |
| --- | --- | --- | --- | --- | --- | --- |
| Initial | 1.0000 | 0.000 | 1.952 | 8.4 | 0.0 | 91.6 |
| MODEL 1 | 0.9752 | 0.331 | 1.571 | 8.7 | 0.9 | 97.5 |

**Figure 5.**
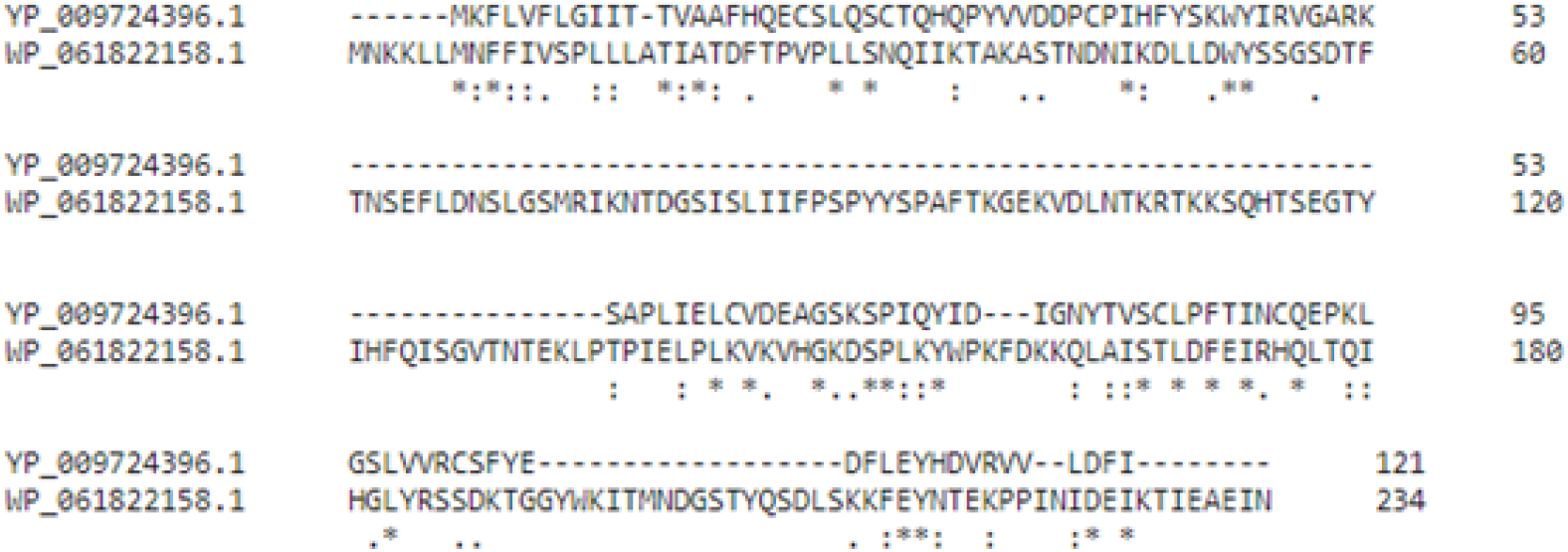
Screenshot of alignment analysis between ORF8 (YP_009724396.1) and TSST (WP_061822158.1) by Clustal Omega. Asterisk: perfect alignment; colon: strong similarity; period: weak similarity.

**Figure 6.**
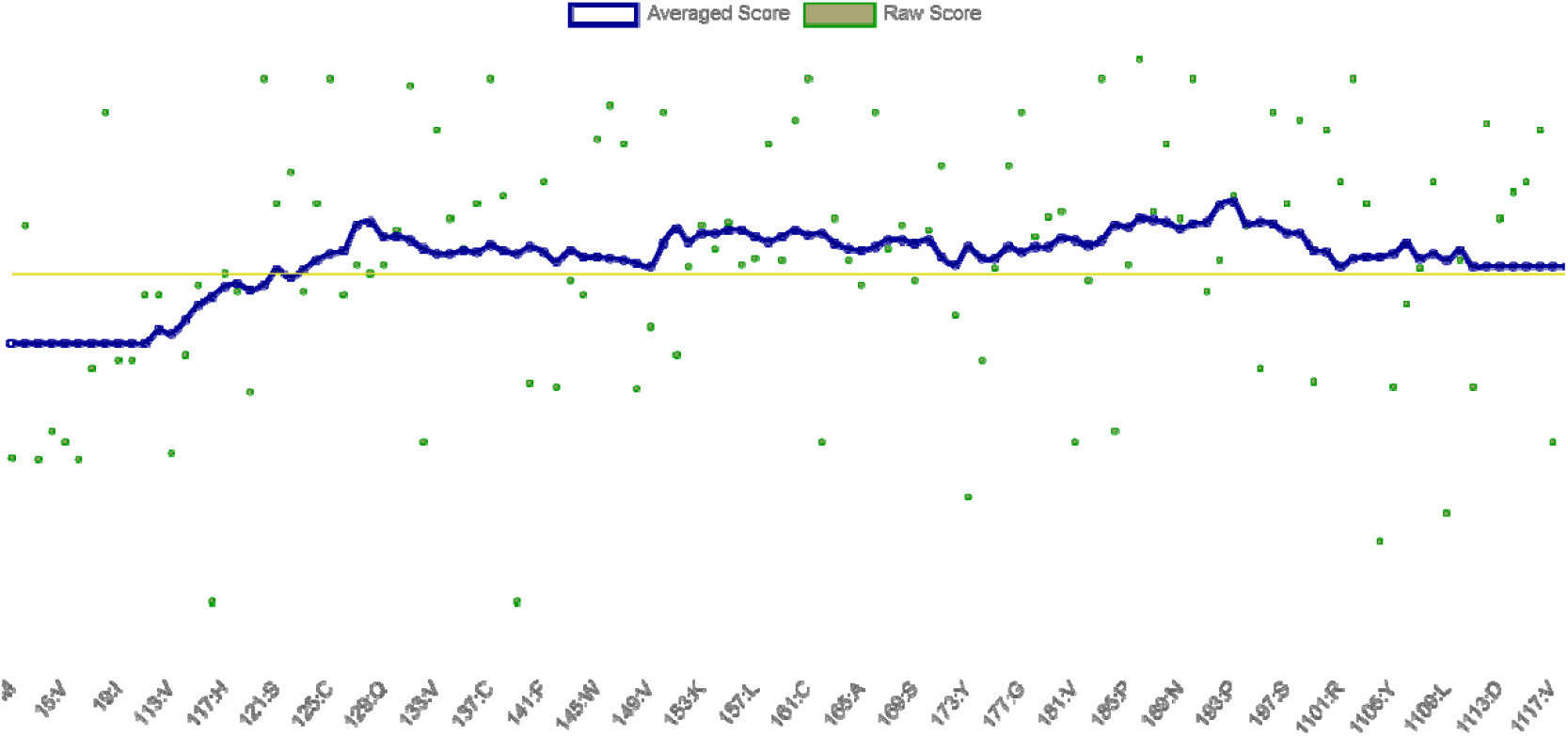
Verification of the correctness of the models according to VERIFY-3D for or monomeric ORF8. 82.64% residues were above 0.2.

**Figure 7.**
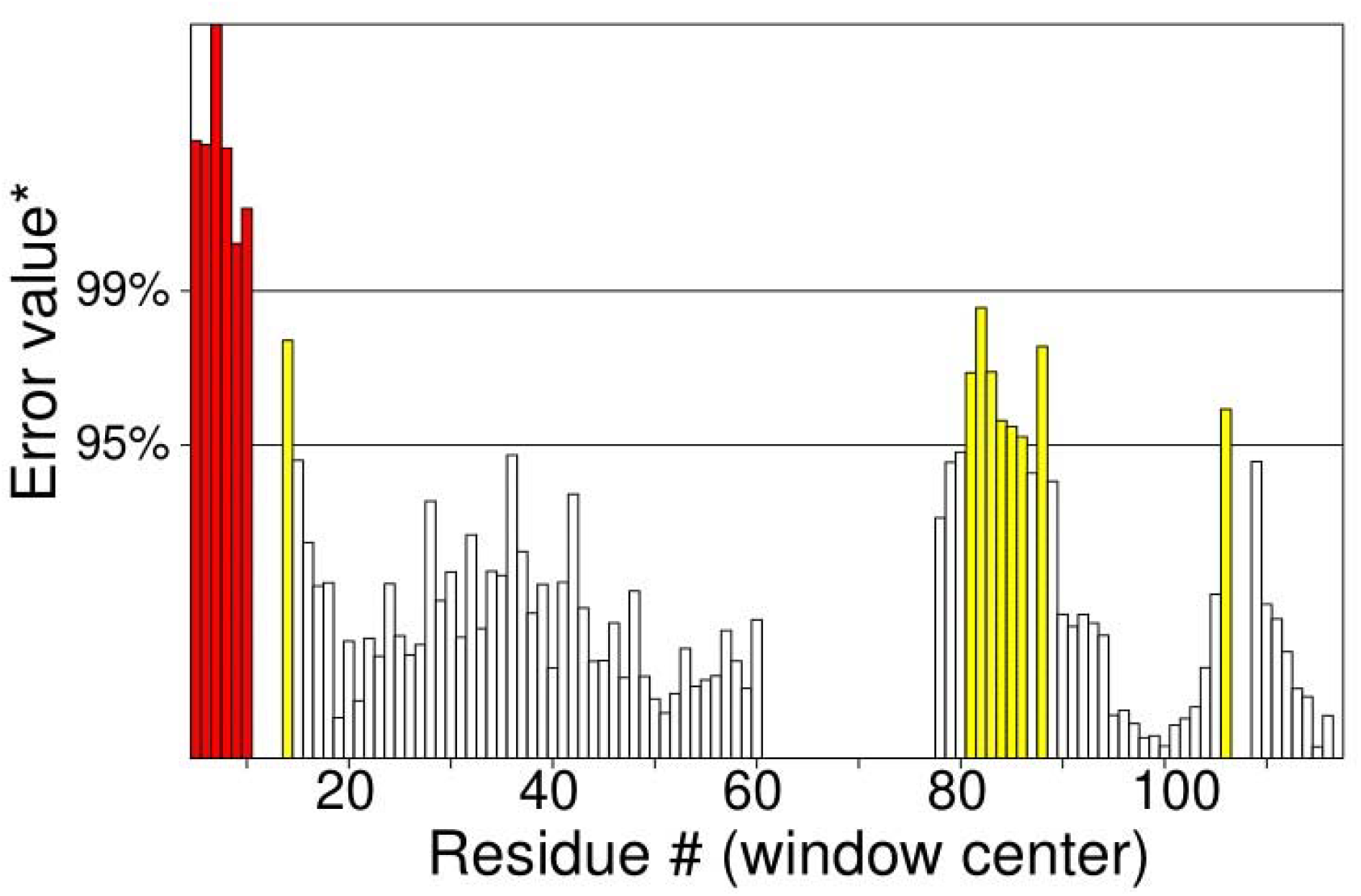
Errat plot for ORF8. Quality factor: 83.333.

Finally, Ramachandran plot analyses were carried out by Pro Check. (Figure 8). Regarding ORF8, the results showed that 92.5% residues were in the favored region, 7.5% were in allowed region and 0% in disallowed region. The results for quality of monomeric ORF8 by different model evaluation tools were summarized in Table 4. Overall, the predicted model quality was considered as good and then was used for further analysis.

**Table 4.** Validation scores of the ORF8 from different model verification tools.

| Verify 3D | ERRAT | Ramachandran analysis |  |  |
| --- | --- | --- | --- | --- |
|  |  | Favored region | Additional<br>Allowed region | Disallowed<br>region |
| 82.64% | 83.3% | 92.5% | 7.5% | 0% |

**Figure 8.**
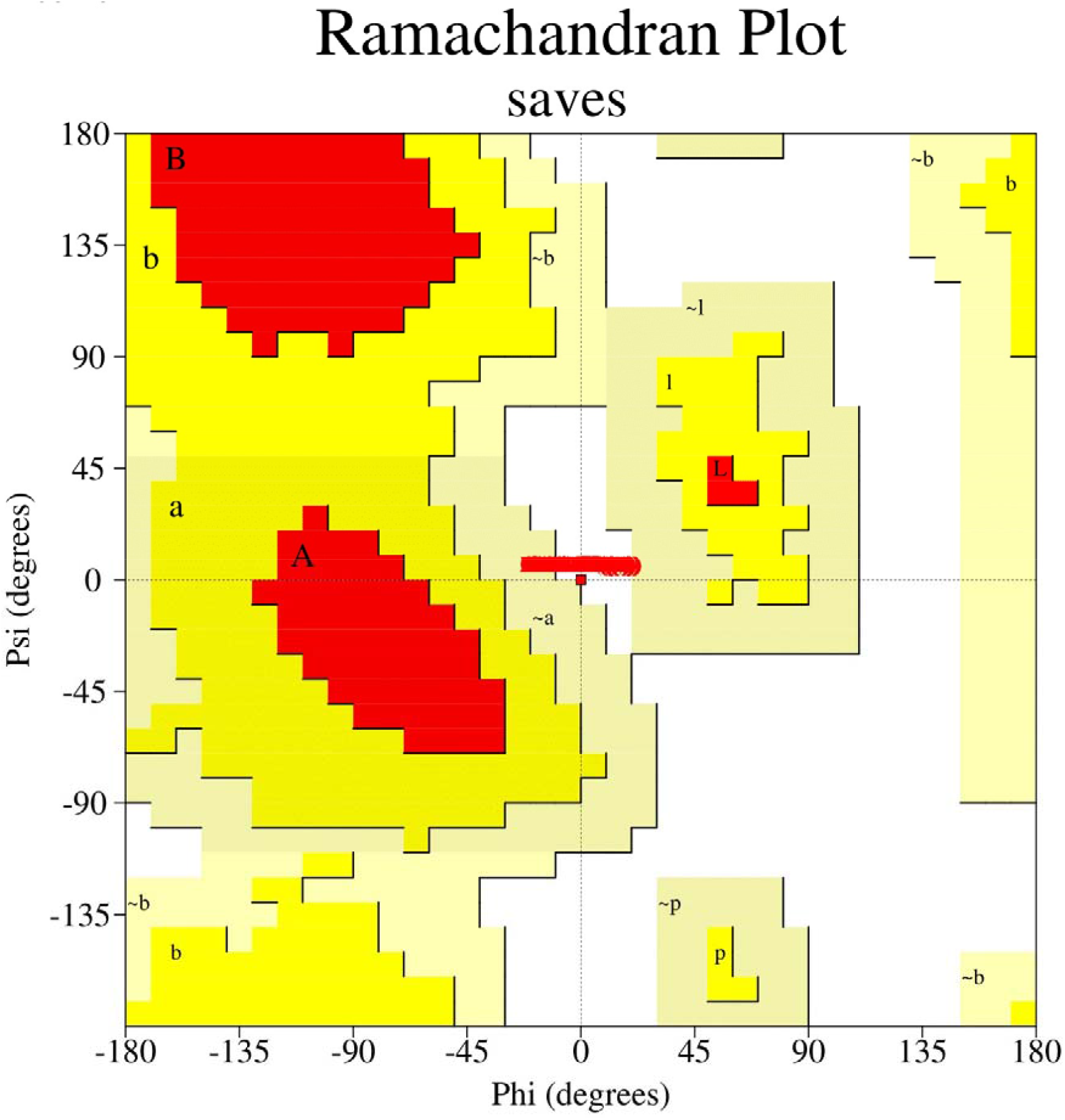
Ramachandran plot of Monomeric ORF8 obtained by PROCHECK, 92.5% residues in favorable regions; 7.5% residues in additional residue regions; 0.0% residues in generously regions; 0.0% residues in disallowed regions.

### 7. Binding site residues prediction

To carry out the docking, binding sites residues were predicted for monomeric ORF8, homodimeric ORF8, Vβ 2.1 and TRBV11-2. The analysis by META-PPISP and SPPIDER of ORF8 showed that monomeric and dimeric ORF8 models did not share any common amino acid in the predicted binding region. In the monomeric model, binding site spanned the center of the molecule, from aa 35 to 43, and the C-terminal region, from aa 104 to 112. In contrast, in the dimeric model, the binding site spanned nears the N-terminal, from aa 22 to 27, and near the C-terminal, from aa 85 to 92. Notably, both the region spanning aa 35 to 43 in the monomeric model and the region spanning aa 85 to 92 in the dimeric model overlapped with the predicted ORF8 similitude region to TSST, according to the BLAST analysis (Figure 3). These regions therefore may bear superantigenic properties. Regarding Vβ chains, the analysis showed that for each Vβ chain, predicted active residues were distributed across the entire molecule. We took the list of common residues from the predicted residues for each protein by SPPIDER and Meta-PPISP and used them as an input parameter for docking, as summarized in table 5.

**Table 5.** List of binding site residues for monomeric ORF8, homodimeric ORF8, Vβ 2.1 and TRBV11-2

| Proteins | Tools | Chains | Binding residues |
| --- | --- | --- | --- |
| Monomeric ORF8 | SPPIDER | A | 1,2,3,4,5,6,7,9,10,11,12,13,14,15,16,17,18,19,20,22,26,30,32,34,35,38,39,40,41,42,43,45,49,52,54,56,57,59,61,62,64,65,66,68,69,70,71,72,73,74,75,76,77,78,85,101,102,104,105,106,107,108,109,110,111,112,113,115,116,117,118,119,120,121 |
|  | METPPISP | A | 4,6,7,17,34,35,38,39,40,41,42,43,44,104,105,106,107,108,109,110,112 |
|  | Common residues | A | 4,6,7,35,39,40,41,42,43,104,105,106,107,108,109,110,112 |
| Homodimer ORF8 | SPPIDER | A | 1, 2, 5, 7, 9, 13, 15, 17, 18, 19, 20, 21, 22, 23, 24, 25, 26, 27, 28, 32, 36, 37, 39, 40, 41, 42, 43, 44, 45, 46, 47, 48, 49, 50, 51, 52, 53, 54, 55, 56, 57, 58, 59, 60, 61, 62, 63, 64, 65, 66, 70, 75, 76, 82, 85, 86, 87, 88, 89, 90, 91, 92, 93, 94, 101, 102 |
|  |  | B | 1, 2, 5, 7, 9, 12, 13, 15, 16, 17, 18, 19, 20, 21, 22, 23, 24, 25, 26, 27, 28, 29, 32, 33, 35, 36, 37, 38, 39, 40, 41, 42, 43, 44, 45, 46, 47, 48, 49, 50, 51, 52, 53, 54, 55, 56, 57, 58, 59, 60, 61, 62, 63, 64, 65, 69, 74, 75, 81, 84, 85, 86, 87, 88, 89, 90, 91, 92, 93, 97, 100, 101 |
|  | METPPIS<br>P | A | 18 21 22 23 24 25 26 27 82 85 86 87 88 89 90 91 92 93 |
|  |  | B | 18 19 20 21 22 23 24 25 26 27 28 81 84 85 86 87 88 89 90 91 92 |
|  | Common<br>residues | A | 18 21 22 23 24 25 26 27 82 85 86 87 88 89 90 91 92 93 |
|  |  | B | 18 19 20 21 22 23 24 25 26 27 28 81 84 85 86 87 88 89 90 91 92 |
| Vβ 2.1 | SPPIDER | B | 1,2,4,6,7,8,9,11,25,26,37,38,39,40,41,42,43,44,45,85,89,90,92,99,101,102,103,104,105,106,107,108,109,110,111,113,115,117 |
|  | METPPIS<br>P | B | 2,4,7,8,9,33,35,36,37,38,42,43,44,45,89,90,92,106,107,108,109,110,111,113 |
|  | Common<br>residues | B | 2,4,7,8,9,37,38,42,43,44,45,89,90,92,106,107,108,109,110,111,113 |
| TRBV11-2 | SPPIDER | E | 3,4,6,7,28, 29, 30, 38, 40, 41, 43, 44, 46, 52, 53, 97, 99, 100, 101, 102, 103, 104, 105, 106, 107, 108, 109, 110, 112, 129, 130, 131, 132, 133, 149, 175, 179, 241, 242, 245, 248, 249 |
|  | METPPIS<br>P | E | 3, 4, 27, 28, 29, 30, 31, 51, 52, 53, 54, 55, 97, 98, 99, 100, 101, 102, 103, 104, 105, 106, 107, 108, 109, 110 |
|  | Common<br>residues | E | 3,4,28,29,30,52,53,97,99,100,101,102,103,104,105,106,107,108,109,110 |

### 8. Docking

Docking of monomeric and homodimeric ORF8 with Vβ 2.1 and TRBV11-2 were carried out by ClusPro 2.0 webserver by providing the PDB’s of the models along with binding site residues. Different clusters were generated by ClusPro 2.0. The best cluster was selected based on maximum numbers of members and more negative energy score. The score for different docked complexes were summarized in table 6.

**Table 6.** Docking score for monomeric ORF8 and homodimeric ORF8 with Vβ 2.1 and TRBV11-2

| Complex | Cluster | Members | Representative | Weighted Score |
| --- | --- | --- | --- | --- |
| Monomeric ORF8 with Vβ 2.1 | 0 | 162 | Center | -1949.1 |
|  |  |  | Lowest Energy | -2070.1 |
| Monomeric ORF8 with TRBV11-2 | 0 | 248 | Center | -1811.3 |
|  |  |  | Lowest Energy | -1855.2 |
| ORF8 homodimer with TRBV11-2 | 0 | 66 | Center | -2409.4 |
|  |  |  | Lowest Energy | -2409.4 |
| ORF8 homodimer with Vβ 2.1 | 0 | 120 | Center | -2023.9 |
|  |  |  | Lowest Energy | -2097.4 |

### 9. ORF8 and Vβ chains interactions

Analysis of interacting residues between monomeric ORF8 and homodimer ORF8 with Vβ 2.1 and TRBV11-2 were carried out in PyMol version 2.4.1.

#### 9.1 Monomeric ORF8-Vβ 2.1

We found that monomeric ORF8 interacted with Vβ 2.1 through 6 aa and formed 10 H bonds with 8 aa of the Vβ 2.1 chain. (Figure 9a) The residues Ser 9, Arg 114, Arg 114, Arg 114, Gln 38, Gln 38, Thr 106, Val 4 and Phe 109 of monomeric ORF8 formed H bonds with Ala 14, Leu 4, Gly 8, Leu 7, Cys 37, Ile 39, Tyr 105, Glu 110 And Glu 110 of Vβ 2.1.

**Figure 9.**
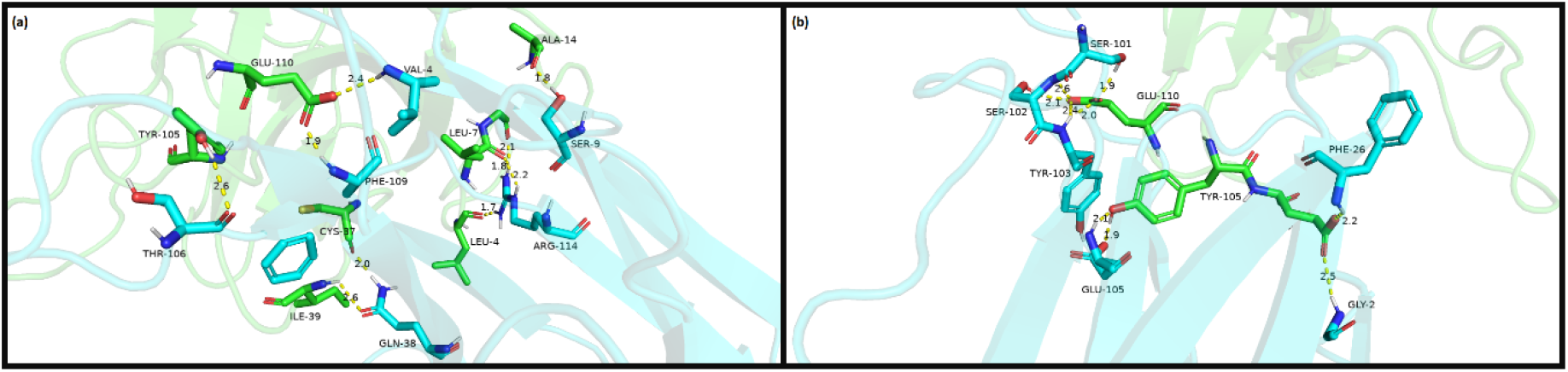
Interacting residues by H bonds between monomeric ORF8 with a) Vβ 2.1 and b) TRBV11-2. Color residues: blue is ORF8; green is Vβ chains.

#### 9.2 Monomeric ORF8-TRBV11-2

Monomeric ORF8 interacted with TRBV11-2 through 6 aa and formed 9 H bonds with 3 aa of the TRBV11-2 chain (figure 9b). The residues Gly 2, Phe 26, Ser 101, Ser 102, Tyr 103 and Glu 105 of monomeric ORF8 formed hydrogen bonds with Glu 106, Glu 106, Glu 110, Glu 110, Glu 110 and Tyr 105 of TRBV11-2.

The amino acids that formed H bonds between monomeric ORF8 with Vβ 2.1 and TRBV11-2 are summarized in table 7.

**Table 7.**
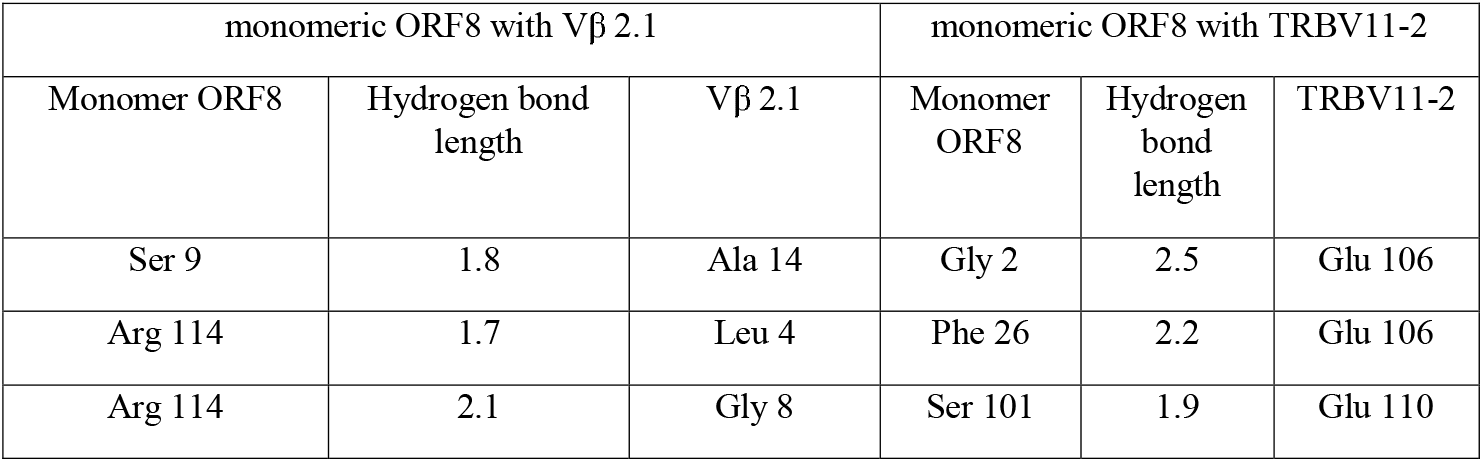
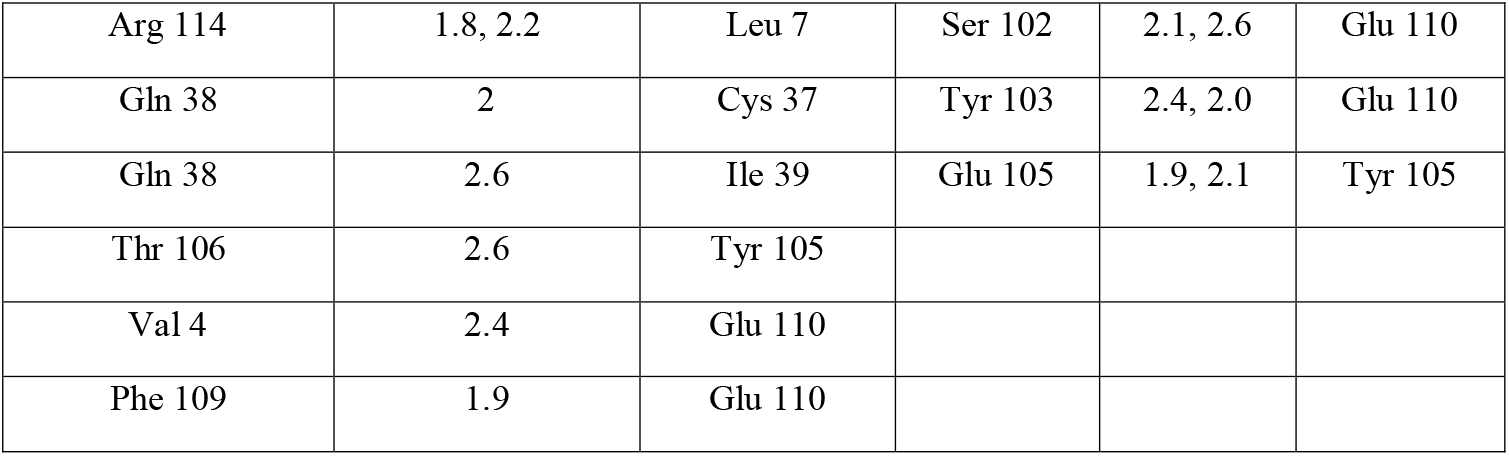
List of interacting residues along with hydrogen bonds length between monomeric ORF8 with Vβ 2.1 and TRBV11-2

| monomeric ORF8 with Vβ 2.1 | | | monomeric ORF8 with TRBV11-2 | | |
| --- | --- | --- | --- | --- | --- |
| Monomer ORF8 | Hydrogen bond length | Vβ 2.1 | Monomer ORF8 | Hydrogen bond length | TRBV11-2 |
| Ser 9 | 1.8 | Ala 14 | Gly 2 | 2.5 | Glu 106 |
| Arg 114 | 1.7 | Leu 4 | Phe 26 | 2.2 | Glu 106 |
| Arg 114 | 2.1 | Gly 8 | Ser 101 | 1.9 | Glu 110 |
| Arg 114 | 1.8, 2.2 | Leu 7 | Ser 102 | 2.1, 2.6 | Glu 110 |
| Gln 38 | 2 | Cys 37 | Tyr 103 | 2.4, 2.0 | Glu 110 |
| Gln 38 | 2.6 | Ile 39 | Glu 105 | 1.9, 2.1 | Tyr 105 |
| Thr 106 | 2.6 | Tyr 105 |  |  |  |
| Val 4 | 2.4 | Glu 110 |  |  |  |
| Phe 109 | 1.9 | Glu 110 |  |  |  |

#### 9.3 Homodimer ORF8-Vβ 2.1

Homodimeric ORF8 interacted with Vβ 2.1 through its two chains (Figure 10a). Chain A formed 1 H bond and Chain B formed 8 H bonds with Vβ 2.1. The ser 7 of Chain A formed a H bond with Asp 27 of Vβ 2.1. Similarly, the residues Ile 22, Tyr 85, Glu 90, Tyr 91, Leu 89, Asp 93, and Lys 36 of chain B formed H bonds with Gln 38, Tyr 36, Gln 107, Gln 107, Thr 106, Ala 2, Asp 27 of Vβ 2.1.

**Figure 10.**
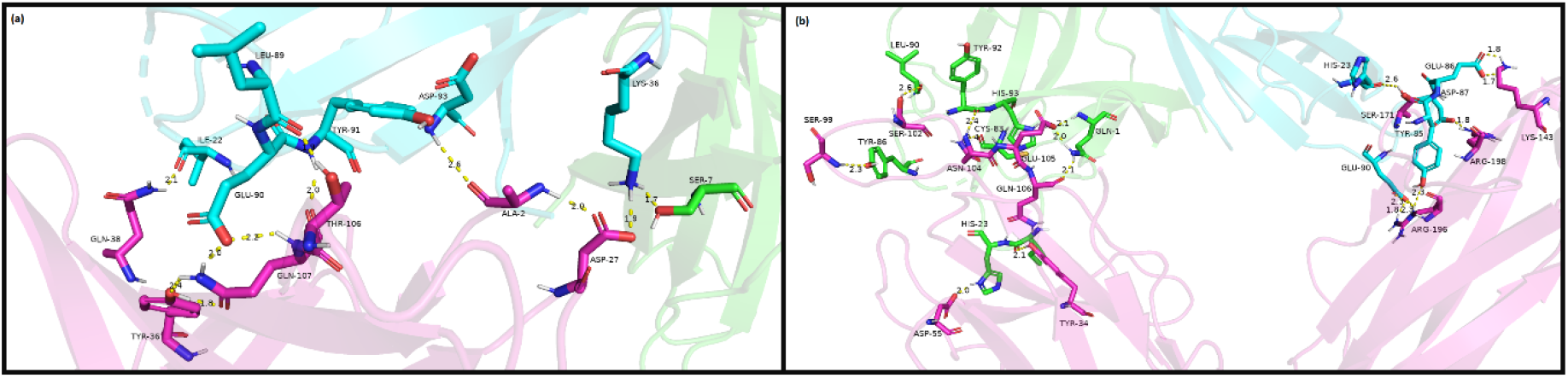
The amino acids making hydrogen bonds between homodimeric ORF8 with Vβ 2.1 and TRBV11-2. Color residues: blue is chain B ORF8; green is chain A ORF8; pink is TRBV11-2 chain.

#### 9.4 Homodimer ORF8-Vβ 2.1

Homodimeric ORF8 interacted with TRBV11-2 through its two chains (Figure 10b). Chain A of homodimer formed 10 H bonds and chain B formed 8 H bonds with TRBV11-2 The residues Tyr 86, His 23, Ile 22, Leu 90, Tyr 92, Lys 83, His 93, Gln 1 and Gln 1 of chain A formed H bonds with Ser 99, Asp 55, Tyr 34, Ser 102, Asn 104, Asn 104, Glu 105, Glu 105 and Gln 106 of TRBV11-2. Similarly, the residues His 23, Glu 86, Asp 87, Tyr 85 and Glu 90 of chain B of homodimer ORF8 formed H bonds with Ser 171, Lys 143, Arg 198, Arg 196 and Arg 196. The amino acids that formed H bonds between homodimer ORF8 with Vβ 2.1 and TRBV11-2 are summarized in table 8. This analysis shows that complexes formed interactions with different H bonds number. In descending order on the bases of maximum to minimum H bonds, we had homodimer ORF8–TRBV11-2 (18 H bonds), monomeric ORF8–Vβ 2.1 (10 H bonds), homodimer ORF8–Vβ 2.1 (9 H bonds), and monomer ORF8–TRBV11-2 (9 H bonds). Therefore, homodimer ORF8 formed the maximum number of H bonds and thus could be the most stable complex.

**Table 8.** List of interacting residues along with hydrogen bonds length between homodimer ORF8 with Vβ 2.1 and TRBV11-2

| Homodimer with Vβ 2.1 | | | | | | Homodimer with TRBV11-2 | | | | | |
| --- | --- | --- | --- | --- | --- | --- | --- | --- | --- | --- | --- |
| Vβ 2.1 | Length | Chain b of ORF8 | Vβ 2.1 | Length | Chain A ORF-8 | TRBV11-2 | Length | Chain B ORF8 | TRB V11 -2 | Length | Chain A ORF8 |
| Gln 38 | 2.1 | Ile 22 | Asp 27 | 2 | Ser 7 | Ser 171 | 2.6 | His 23 | Ser 99 | 2.3 | Tyr 86 |
| Tyr 36 | 1.9 | Tyr 85 |  |  |  | Lys 143 | 1.8,1.7 | Glu 86 | Asp 55 | 2 | His 23 |
| Gln 107 | 2.0, 2.2 | Glu 90 |  |  |  | Arg 198 | 1.8 | Asp 87 | Tyr 34 | 2.1 | Ile 22 |
| Gln 107 | 2 | Tyr 91 |  |  |  | Arg 196 | 2.3 | Tyr 85 | Ser 102 | 2.6 | Leu 90 |
| Thr 106 | 1.9 | Leu 89 |  |  |  | Agr 196 | 2.3,2.3, 1.8 | Glu 90 | Asn 104 | 2.4 | Tyr 92 |
| Ala 2 | 2.6 | Asp 93 |  |  |  |  |  |  | Asn 104 | 2.4 | Lys 83 |
| Asp 27 | 1.9 | Lys 36 |  |  |  |  |  |  | Glu 105 | 2 | His 93 |
|  |  |  |  |  |  |  |  |  | Glu 105 | 2.1,2.0 | Gln 1 |
|  |  |  |  |  |  |  |  |  | Gln 106 | 2.1 | Gln 1 |

### 10. Binding affinity

Binding affinity of the complexes was analyzed using Prodigy. The analysis showed that all forms of ORF8 were predicted to strongly bind Vβ chains (Table 9). We found that homodimer ORF8 had lower Kd than monomer ORF8 and then higher affinity. In both cases, the values were indicative of a strong interaction between viral and immune proteins. Notably, such values were similar to that predicted for Sag toxin-Vβ 2.1 complex. Thus, different forms of ORF8 may have an impact on Vβ chain-associated immune response. The analysis of ΔG showed that the complexes of monomeric and homodimeric ORF8 with TRBV11-2 had similar binding affinity, while the monomeric ORF8 – TRBV11-2 complex had the lowest binding affinity. In ascending order on the bases of more to less negative ΔG values, we had homodimer ORF8 – TRBV11-2 (−12.9 kcal mol-1), monomeric ORF8 – Vβ 2.1 (−12.5 kcal mol-1), homodimer ORF8 – Vβ 2.1 (−11.8 kcal mol-1), and monomer ORF8 – TRBV11-2 complex (−7.5 kcal mol-1). As a comparison, the ΔG for *S. aureus* – TCR was of -10.4 kcal mol-1, indicating that ORF8, particularly in its homodimeric form had strong binding affinity for TCR Vβ chains.

**Table 9.**
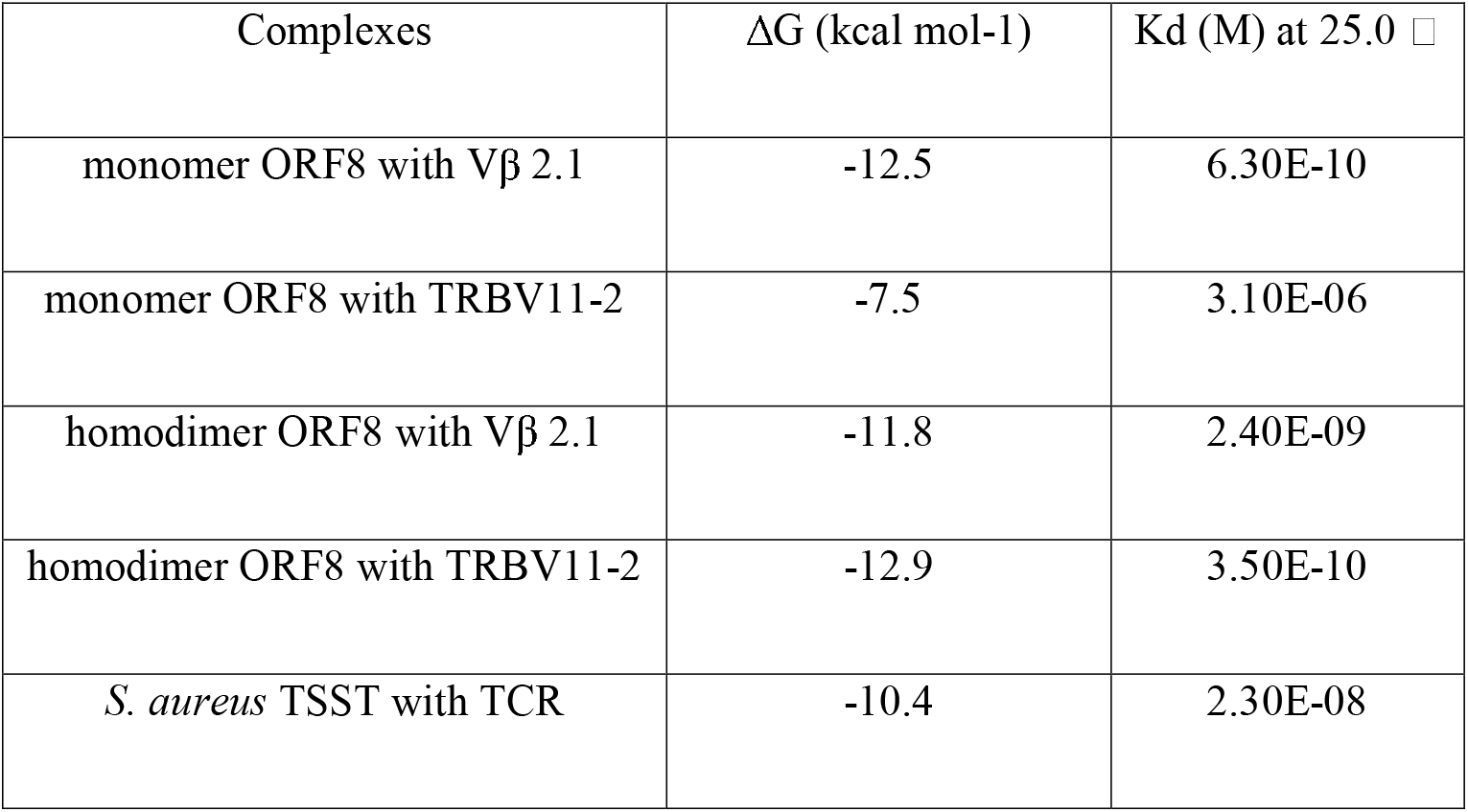
Predicted binding affinity of analyzed complexes expressed in the form of Gibbs free energy and dissociation constant.

## DISCUSSION

The role of ORF8 from COVID-19 in viral spread or disease has not been properly determined. We evaluate the properties of this viral protein as a superantigen. We carry out *in silico* predictive analysis of cellular localization, similarity analysis with TSST and docking prediction with TCR Vβ chain TRBV11-2. The finding that ORF8 contains attributes related entirely to an extracellular and soluble protein, according to the localization analysis, along with the identification of a signal peptide at N-terminal region, clearly indicates that ORF8 may be a circulating protein in blood. This can explain the frequent detection of anti-ORF8 antibodies in sera from COVID-19 patients during all stages of the disease [45]. Experimental evidence of functional activity of this protein so far comes from its intracellular activity. In a study, intracellular ORF8 downregulates MHC-I from *in vitro* cultured cells, thus prevents its elimination by cytotoxic cells. This is a property shared with Nef protein from HIV-1. Like ORF8, Nef is a viral nonstructural protein with the ability to intracellularly interact with, and downregulate MHC-I [46]. Remarkably, Nef is also detected as circulating protein in serum from patients, in concentrations surpassing those from HIV-1 structural proteins by several orders of magnitude [47, 48]. Therefore, similar to Nef, ORF8 might have a dual functional localization, at both intracellular and extracellular milieu. Surprisingly, during the submission process of this work, a paper was published demonstrating that ORF8 actually is a blood circulating protein and its associated to COVID-19 severity [49]. Therefore, the potential role of this protein as a Sag cannot be overlooked.

The striking degree of aa similarity between ORF8 and TSST, according to BLAST and Clustal Omega analysis, provide support to our hypothesis that this viral protein bears Sag determinants. Moreover, the finding that ORF8 aa sequence matches with some aa of TSST considered as essential to Vβ 2.1 docking, suggest that this viral protein might dock Vβ chains by a related mechanism. In addition to ORF8, another *in silico* study has also identified a Sag determinant of 11 aa length in the Spike protein [14]. This Sag has marked similarity to a region of staphylococcal Sag enterotoxin B and may bind directly to Vβ chain of TCR. Further studies will allow to determine the contribution of this spike Sag and ORF8 to MIS and to other forms of systemic thromboinflammation.

Our docking analysis indicates that ORF8 may form a strong interaction with Vβ chains 2.1 and 21 of human TCR through multiple H bonds, which provide stability to protein-protein complexes [50]. The degree of H bonds varies between monomeric and dimeric ORF8. The complex ORF8-TRBV11-2 contain the largest number, and consequently form a more stable interaction and thus may have a deeper effect in T cell function than the other complexes. Moreover, it is worth noting that ΔG and Kd values from ORF8-Vβ chains docking are similar to those from *S. aureus*-Vβ 2.1 docking. Thus, the physical and energetic features of ORF8-Vβ chains docking are comparable to those observed in Sags of clinical relevance. ORF8 also dock with IL-17 receptor and induces the release of IL-17, along with other proinflammatory cytokines in cultured cells and contributes to cytokine storm in a mice model [23]. Then, ORF8 displays a variety of protein-protein interactions that impact the inflammatory response independently of viral replication.

Evidence of ORF8 contribution to the clinical status comes from a study of a group of patients infected with a viral variant carrying a 382 nucleotide deletion in ORF8, thus probably rendering a non-functional protein (51). These patients exhibited lower levels of inflammatory factors along with milder disease at overall and significantly lower oxygen requirements that wild type infected patients. This study, albeit limited, clearly indicates that ORF8 have an impact in disease severity.

In summary, our research establishes that ORF8 contains superantigenic features. Further research will improve our understanding of its role in pathogenesis in order to conceive novel treatments.

## AUTHOR CONTRIBUTION

Gomez-Icazbalceta conceived the hypothesis, designed the research route, performed *in silico* and data analyses, wrote results, figures and manuscript; Hussain performed advanced *in silico* analyses, wrote methods, results, references and figures and analyzed data; Velez-Alavez wrote methods, analyzed data, contributed to discussion and overall review of the manuscript. All authors read and approved the manuscript.

